# Do not let the beginning trap you! On inhibition, associative creative chains, and Hopfield neural networks

**DOI:** 10.1101/2023.09.22.559049

**Authors:** Ronald Mtenga, Mathias Bode, Radwa Khalil

**Affiliations:** School of Electrical and Computer Engineering, Constructor University, Bremen, Germany; School of Business, Social, and Decision Sciences, Constructor University, Bremen, Germany

**Keywords:** Creative Thinking, Hopfield Neural Networks, Patterns, Memory, Associative Chains, Inhibition

## Abstract

Creative thinking stems from the cognitive process that fosters new ideas and problem-solving solutions. Creative cognition in emerging artificial intelligence systems and neural models may reduce complexity in understanding creative cognition. Hopfield Neural Networks (HNN) is a simple neural model known for its biological plausibility to store and retrieve neuron patterns. The primary objective is to demonstrate that ideas, symbolized as patterns of ones and zeros representing clusters of neurons that synchronize their firing, can be stored within HNN and establish connections through correlation. The network can converge towards these ideas by manually adjusting specific state parameters, effectively controlling the overall network activity. When the second closest stored pattern deviated significantly from the input pattern, the network’s ability to converge decreased, enabling it to connect the input and patterns. Thus, we suggest employing HNN for the first time to create a model that emulates creative thinking processes, including making meaningful links between seemingly unrelated ideas.

We implemented certain modifications to the original HNN, including introducing pattern weight control, which provides a robust representation for content addressable memory and illustrates conceptual links in stored data, a step towards the larger framework of creativity. We have made progress in identifying two mechanisms that could assist in managing the dynamics of the network and the formation of associative links. These mechanisms are related to the activation threshold of the neurons and the inhibitory stimulus on the stored patterns.

## Introduction

Finke et al. (1996) conducted a study in which they reported that individuals with creative abilities can engage in a more expansive form of thinking that involves making connections between ideas and concepts. This cognitive process enables them to divert their attention from familiar tools and instead focus on exploring the relationships between various elements. This approach facilitates the identification of less obvious similarities, hence enabling the emergence of insights that would otherwise be unachievable if one’s focus were exclusively on the surface-level characteristics of the task (Beaty et al., 2023; Kenett & Beaty, 2023; Luchini et al., 2023).

There is a consensus among scholars that creative products should effectively convey original, valuable, and potentially surprising concepts (Corazza, 2016; Gardner, 1994; Runco & Jaeger, 2012; Torrance, 1974; Wilson et al., 1954). As a result, it is widely acknowledged that creative thinking is esteemed for its capacity to produce answers that are novel, useful, and surprising (Han et al., 2021; Khalil & Moustafa, 2022; Mastria et al., 2022). However, creativity is a complex concept that can manifest itself in a multitude of ways, presenting a significant challenge in defining its precise nature (Abraham, 2013; Gaut, 2010; Kaufman & Beghetto, 2009; Sawyer, 2006, 2011; Squalli & Wilson, 2014; Tardif & Sternberg, 1988). One potential approach to mitigate the intricacy and redundancy is to focus on the neural network structures and their associated learning rules, as Khalil and Moustafa proposed (2022).

Therefore, we aim to integrate this creative cognitive process-based semantic association (Beaty & Kenett, 2023; Kenett & Beaty, 2023; Li et al., 2021) within a conceptual framework designed for a creative and feasible neural network system utilizing a Hopfield neural network (HNN). HNN, a mathematical tool invented by John J. Hopfield in 1982, is widely acknowledged for its capacity to store and retrieve information (McEliece et al., 1987). Establishing the HNN model was a significant achievement in neural computation and modeling (Abe, 1989, 1993; Hopfield, 1982; KÖksal & Sivasundaram, 1993; Ramsauer et al., 2021).

HNN is a recurrent associative neural network model with the distinctive feature of receiving input from all neurons in the network. This unique feature enables the HNN to effectively recover memories even when presented with incomplete or noisy input, as described initially by Hopfield in 1982. The HNN employs iterative updating to synchronously or asynchronously update the states of its neurons to converge towards one of the stored memories and attain a stable state that recalls the information it was trained on (Krotov & Hopfield, 2016). Therefore, the utilization of HNN has been observed in tasks related to associative memory, such as the association of input with its most similar stored pattern or the identification of unknown systems through the computation of optimum coefficients by the learning mechanism to effectively model the nonlinear system (McEliece et al., 1987; Wang et al., 2003, 2005).

We chose the HNN model to develop a semantic association-based creative conceptual framework because it recalls patterns by considering the correlation between neurons using the weights matrix, not storing them (Beaty et al., 2023; Kenett & Beaty, 2023; Luchini et al., 2023). Furthermore, the topology of HNN closely mirrors human memory, which encodes many neurons that fire together in response to “microfeatures” (Gabora, 2010). Creative thinking allows individuals to switch between analytical and associative modes to find solutions (Dietrich, 2004b, 2004a; Finke, 1996; Xie et al., 2021), and thus the act of creativity enables individuals to view the environment in novel ways, identify concealed patterns, and establish associations between seemingly disparate occurrences, resulting in the production of exceptional outcomes (Abraham, 2018; Herrmann, 1981; Khalil & Demarin, 2023; Khalil & Moustafa, 2022). The model proposed by Mendelson in 1974 utilized defocused and focused attention to explicate the semantic hierarchy of creative associations. The idea of the associative foundation of the creative process aligns with Mednick’s earlier introduction of this term (1962). Gabora (2010) called this context attention, which may affect creative problem-solving, and explained that each neuron saves several items and distributes them among neuron assemblies (Morris, 1999; Rumelhart et al., 1986). This view means that concepts and ideas are widely distributed within neural assemblies, where each neuron is used and re-used in a phenomenon called neural re-entrance (Calvin, 1988).

Neurons in clusters can be approximated as patterns of 1s and 0s, with 1 representing high-rate neurons and 0 representing low-rate neurons; this allows for the manipulation of concepts using HNN. These notions are learned through repeated excitation of the same neurons, which reduces the synapses’ electrical signal resistance and increases the likelihood that they will excite each other when a signal is present. If we consider internal or external stimuli stimulate some of these neurons, the rest of the neurons with the closest associative link will be triggered, causing an avalanche effect that recalls the stored concept.

We describe this semantic linkage as a radio button that we can turn to tune whether the network can solve problems creatively or if it shuts down entirely; the threshold is used as a parameter. We refer to it as the “**first link of creativity”**; in this context, we are invoking the “**second link of creativity”** to facilitate exploring alternative solutions inside our network. By manipulating the button, it is possible to selectively suppress specific patterns, which could be an analog to inhibition, facilitating the creative functioning of the HNN and identifying other patterns with which our input can be linked.

## Methods

This section describes the neural network structure and implementation of HNN.

### Neural Network Structure

The network structure includes patterns and weight matrix updating (Fig. 1).

**Figure 1.**
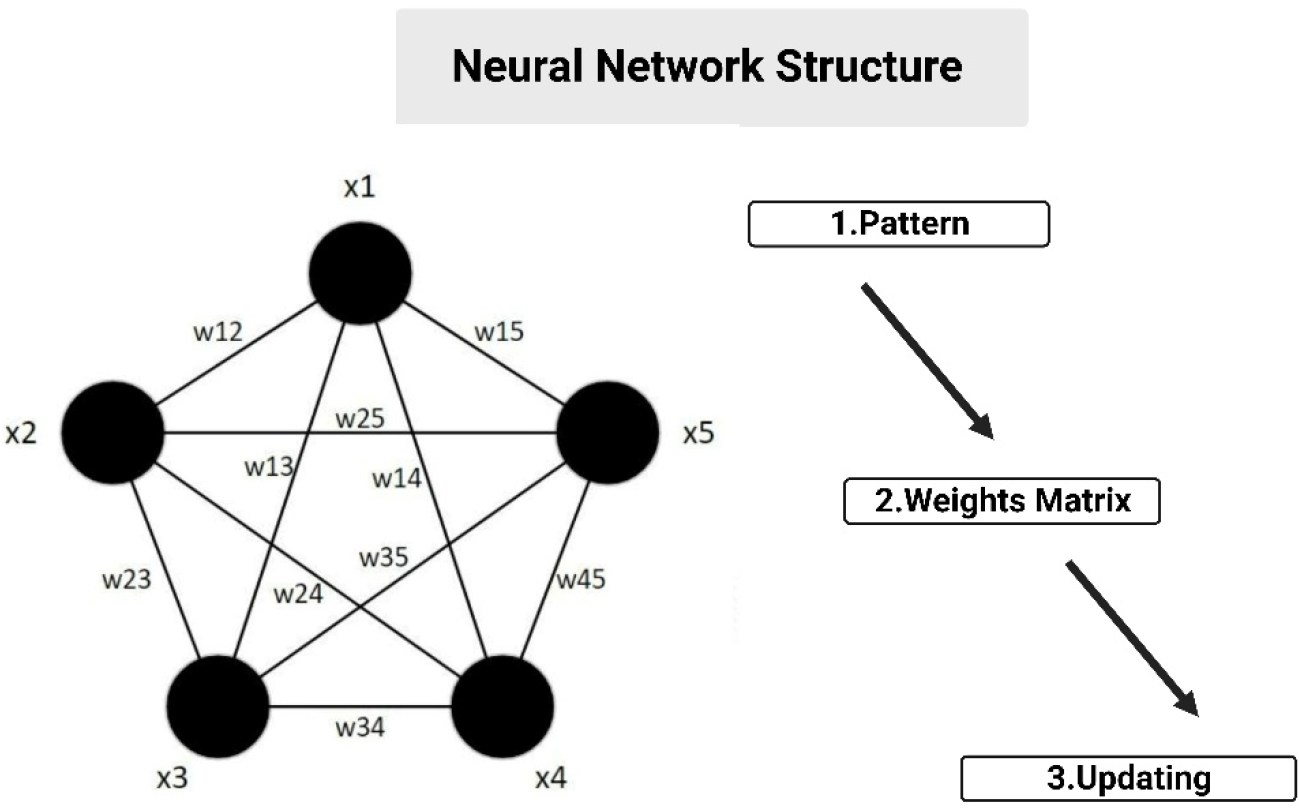
Description of the neural network structure of Hopfield neural networks (HNN). Mutual activation of neuron pairs determines system node weights. If two neurons are active simultaneously, their weight is one; otherwise, it is zero. Each time a new pattern is added to the system, the weights between activated nodes are strengthened.

In 1982, John Hopfield implemented the Hebbian learning rule as the sum of the outer products of the patterns to be stored, creating an auto-correlation matrix that may be normalized by the number of patterns stored (Hopfield, 1982). Each matrix entry represents the average neuron activity relative to another. Setting the auto-correlation matrix diagonal to zero creates a symmetric matrix with average cross-activation values and zero self-activation. An energy function that decreases or remains constant after each neuron update and contains local minima matching to stored patterns may also be utilized to understand pattern storage (Morris, 1999). The Hebb rule refers to increases in the weight between neurons when a pattern is introduced when they are firing together (Klein, 2011; Morris, 1999). When an input pattern contains misaligned information, it is equivalent to “straining” these weights. Once the system can update, the auto-correlations of the closest stored pattern will overwhelm this distorted information, bringing it closer to its activation states.

The original HNN was described as a simple auto-associative memory where the bidirectional weights, *w*_*ij*_ of a fully connected single-layer neural network, are computed by finding the outer product of each of the discrete binary patterns (**x**_k_, …, **x**_M_) that are to be stored, and summing up the resulting matrices:

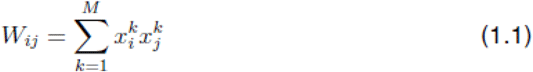

We simulated HNN with N = 1024 neurons with patterns chosen from the binomial distribution, and simulations were performed for varying p (where *p* = *P*(σi = 1)). As a result, these weights are learned iteratively, with a weight’s value only reinforced if the learned pattern activates the same pair of neurons. This process is similar to the learning process in the brain; neurons process sensory and contextual information about an idea fire together as the person experiences it, and with prolonged exposure, synaptic resistance decreases, allowing neurons activated by certain aspects of the concept to drive the activation of every other neuron linked to it (Gage & Hickok, 2005).

According to Gabora et al. (2010), how neurons associated with storing concepts are distributed over a cell assembly plays a critical role in the brain’s capacity to retain memories that can be accessed and facilitate associative thinking. Whether to normalize the weights to limit their levels adequately is arbitrary. However, it does not affect the updating of the system, as the relative magnitudes of the weights matter in this process rather than their absolute values. Once the weights are learned, meaning the patterns are now local minima in the system, the neural system is excited by providing a different input from the stored patterns; it can cool down and settle into a local minimum closest to the input in a process known as simulated annealing.

Given a pattern, **X**_k_ has been learned and is in a stable state in the system:

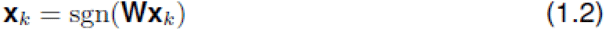

**W** is the weight that also depends on previously learned patterns, and sgn() is the non-linearity that defines our state space of unipolar binary vectors. We used asynchronous updating of the network to analyze all the inputs that feed into the activation of a single neuron in the system. Asynchronous updating provides several advantages over synchronous updating; it allows for faster convergence to stable states and better recall of stored patterns with noise, thus helping avoid oscillations and limit cycles (El Boustani & Destexhe, 2009; Kobayashi, 2021; Moyer et al., 2014). Each neuron update will either decrease the energy of the whole system or keep it constant, which is consistent with Hopfield’s definition of the energy function (Hopfield, 1982):

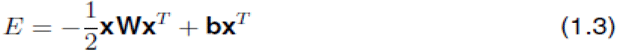

Here b are the bias parameters applied to each network neuron and represent the external influences on that neuron’s activation. We assume the biases are zero; thus, the energy is only given by the first quadratic term. The 1/2 factor is there only to remove the coefficients arising from the quadratic matrix computation, and the negative sign ensures we have a convex quadratic multidimensional energy plane capable of having more than one minimum, unlike a concave quadratic. Our goal is to develop a neurocomputational model for creative thinking based on semantic associations (Beaty et al., 2023; Benedek et al., 2023; Gabora, 2010; Gerver et al., 2023); thus, it is logical to have less correlated concepts to identify the unexplored connections. Based on this reasoning, our patterns represent unrelated concepts, suggesting that a substantial proportion of neurons firing per pattern is unnecessary (i.e., meaning less likelihood for correlation)(Amari & Maginu, 1988; Anishchenko & Treves, 2006; Hopfield, 1982; McEliece et al., 1987).

### 1. Pattern Choice

The choice of patterns to represent the activation of neurons was extensively deliberated. The utilization of binary activations to simulate the high and low activations of neurons was evident from the beginning. Neurons activate at a firing rate and in the continuous range. It has been suggested that bipolar binary representations are favorable where 0 is mapped to −1 and 1 to +1 (Roxin et al., 2008; Šíma et al., 2000):

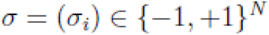

Bipolar activations enable the storage of patterns with both positive and negative values, increasing the network’s representational capacity. This binary mode reflects a simplistic and discrete view of how the neurons act (Šíma et al., 2000). Every neuron within the network is interconnected with all other neurons, with each link possessing a distinct weight analogous to the synapses facilitating communication between neurons. These synaptic weights determine how neurons modify their states in response to this inter-neuronal communication (Ramsauer et al., 2021). The phenomenon of a distinct firing sequence among a collection of individual neurons originating from several regions is commonly known as a recall concept, as described by Hopfield (1982).

Because of the dynamic range of bipolar activations, convergence is less sensitive to noise disturbances. It also simplifies pattern orthogonality; subsequently, bipolarity offers more possibilities for orthogonal vectors than unipolarity. However, we chose unipolar activations because bipolarity does not allow us to activate modestly, as the synaptic contact between neurons determines activation amplitude. For example, bipolarity does not allow us to have low activation since the magnitude of activation is considered in the synaptic interface between neurons. Signals received from surrounding neurons should conquer a potential barrier to activate neurons. A typical neuron has a negative potential relative to its surroundings, but its activation is a question of amplitude more than polarity.

### 2. Weights Matrix

We use a weights matrix *W* to compute the connection of the neurons stored and memorize n patterns in our HNN. We used the Hebbian learning rule for HNN pattern storage, based on the well-known statement, **“Neurons that fire together wire together”** (Klein, 2011; Morris, 1999). Accordingly, if two neurons fire simultaneously, their connection value—the unique weight connecting them—increases, causing their activity to become more correlated over time. (Morris, 1999).

The HNN has the benefit of training in one step by computing the weights matrix W in one operation (Abe, 1989, 1993; Abe & Gee, 1995; Atencia et al., 2005; Joya et al., 2002; KÖksal & Sivasundaram, 1993; Ramsauer et al., 2021). Given their full connectivity, the neurons’ symmetric and bi-directional connections generated W diagonal symmetry. Therefore, the binary input *x* of N neurons is iterated through HNN to get a stable state. A weights matrix *W* facilitates convergence, and the network has been trained and contains all relevant information within *W*; thus, patterns are not required for this operation (Ramsauer et al., 2021; Šíma et al., 2000).

### 3. Updating

To study the effects of the probability of 1s on the dynamics of our network, we consider a network that only contains the associative memory of one pattern. Accordingly, the system receives input from patterns that have been distorted or perturbed, and the system is permitted to update asynchronously for an entire iteration of the network, with each neuron being updated just once. The following equation shows how the j-th neuron of the system is updated asynchronously:

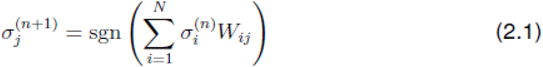

Since σ is a perturbation of the stored pattern, we can further unfold the update equation to show that the total input to the activation of the j-th neuron comes from how close the input is to the stored pattern:

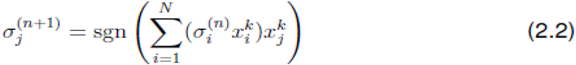

This rearrangement shows that the “closeness” is the inner product of the network input and the stored state. Given the activation function:

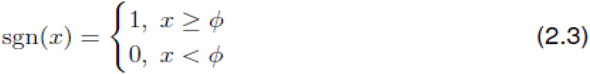

The neuron will be switched on or off if the similarity with the stored pattern is greater or less than a threshold value ϕ. Now, if M patterns are learned instead of one, we can see the effect of the distribution of 1s in the training patterns. Given an input state that is derived from one of the training patterns, we can represent the asynchronous update of the j-th neuron of the system as:

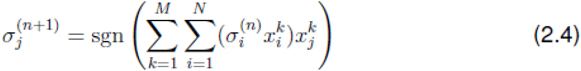

The update has contributions from one pattern and all the learned patterns, showing that the relationship between them is a major factor in whether a neuron will be switched on or off during its asynchronous update. For example, if the input to the system is also one of the learned patterns or is very close to them, we can further expand the updated sum into:

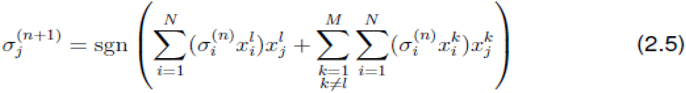

The first term is derived from the outer sum and represents the l-th pattern from M patterns with the largest correlation with the input pattern. Since HNN convergence aims to recover the pattern best resembling the input, this first term represents the input for every j-th neuron update.

The second term represents cross-talk, and in an ideal HNN, it will have slight to no magnitude and thus not interfere with the updating and convergence to the closest pattern. However, the magnitude of the correlation term will be significant, especially once we start increasing the number of patterns being stored. The magnitude of the dot product between the input pattern and the k-th stored pattern and the value of the j-th neuron of the kth pattern indicates whether the j-th neuron will be affected by cross-talk. The correlation determines the dot product value and informs that thresholding will reduce cross-talk if we use uncorrelated learning patterns.

Each k-th learning pattern contributes to cross-talk in the unipolar binary case depending on its 1s and 0s distribution. Cross-talk occurs when the j-th neuron of an interfering pattern is engaged during updating. If the interfering neuron is off, there will be no cross-talk from that pattern. The worst-case scenario is if the update matches the j-th neuron of every other training pattern, causing cross-talk and convergence issues. All interfering patterns will activate the j-th neuron to zero in the best situation; thus, only the lth pattern will be detected.

#### Implementation and Simulation

Python programming language was used to implement the algorithm; the network’s functionality was built using numpy, matplotlib, and random (Davison, 2008). The patterns denoted by ‘n’ were generated by randomly combining zeros and ones, resulting in a cumulative length of N neurons. The code availability is upon request.

### 1. Cross-Talk Effect

To further analyze the cross-talk effect, simulations were performed on different HNNs trained on random patterns with varying p and numbers of training patterns to determine how p affects convergence and capacity.

M = 2 patterns were chosen with length N = 1024 and *p* = *P*(σi = 1) = 0.4.

As a measure of the patterns correlation, the patterns were arranged as the row vectors of a single training matrix, V ∈ 0, 1^(M×N)^, whose squared norm was calculated, resulting in the following symmetric M × M matrix:

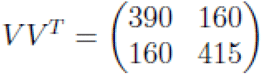

The stability of the training patterns was assessed by feeding a single pattern into the system and observing its iterations. The pattern was considered stable and stored if the system remained in that condition without significant changes. The selection of the activation function threshold, sgn(x), was pivotal during the simulation. The optimal threshold range for the first training pattern was determined to be between 160 and 390, but for the second training pattern, stability was observed within the threshold values ranging from 160 to 415.

The asynchronous update of the jth neuron in the system involves considering the off-diagonal terms in the matrix. These terms represent the cross-talk weight from each pattern when the input pattern is one of the training patterns. Analyzing these weights provides a valuable understanding of the reasons behind the specified threshold ranges.

The j-th neuron will be influenced by the correct pattern and suppress the influence of cross-talk if the threshold is lower than the squared norm of the training pattern used as an input and higher than the dot product with the remaining pattern. If the threshold value is above 390, it will deactivate all neurons. Conversely, if the threshold value falls below 160, excessive cross-talk would occur, leading to instability in the training patterns.

The amount of cross-talk each neuron experiences at each update shows how 1s and 0s affect it. Cross-talk is typically absent, but when it occurs, it is 160 and frequently present (Fig. 2).

**Figure 2.**
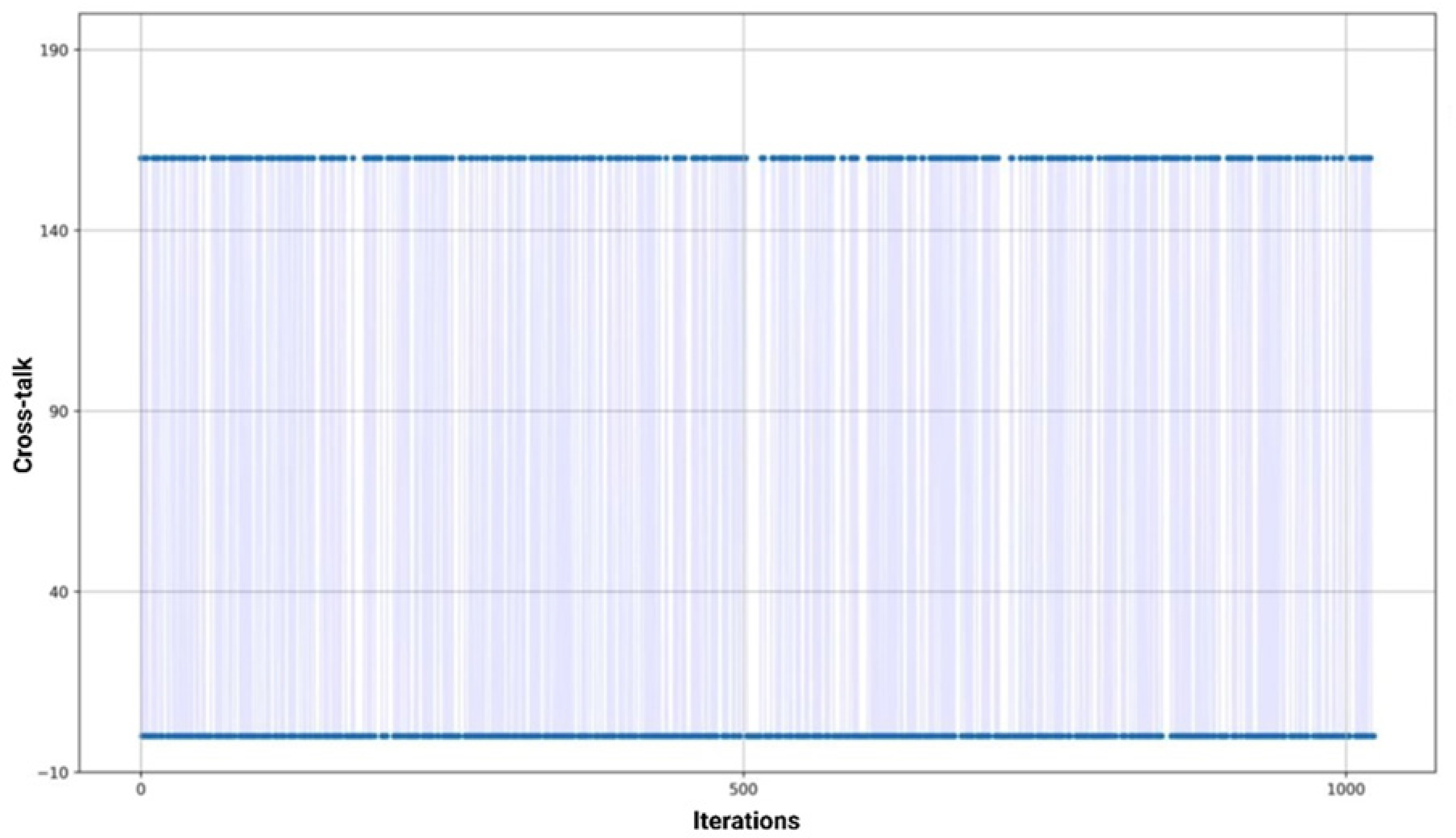
Illustration of the extent of cross-talk encountered by individual neurons at each update, consequently highlighting the influence of binary values (1s and 0s) on their functioning. When cross-talk occurs, it is commonly seen and has a frequency of 160.

When we store more patterns while keeping *p* = 0.4, the total possible cross-talk interferes with retrieving the correct pattern, resulting in spurious stable points or oscillations and limiting cycles in the state space. Spurious states are local minima that the network erroneously stores with the training patterns. They are linear combinations of numerous associated patterns that decrease the network’s storage capacity.

We then simulated M = 4 patterns with the norm square matrix:

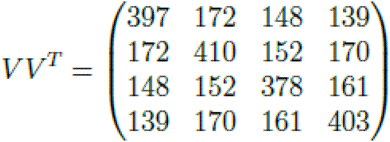

The pivot points represent the closest pattern’s maximum contribution to the input (because we are inputting the patterns), and the remainder of the patterns on a row represents the maximum possible interference. If the first pattern is the input, the worst-case cross-talk is 172 + 148 + 139 = 459, larger than the primary pattern’s contribution, yielding a minimum signal-to-noise ratio (SNR) of 397/459. We simulate the cross-talk over 1024 iterations and select a threshold slightly below the squared norm of our input’s initial training pattern to examine how this affects network stability.

A parameter search was performed to determine the appropriate threshold values. The search considered values within the specified range, which was calculated as 90% of the square norm of the first training pattern to 150% of the square norm. The range was divided into steps of 0.005. The decision to choose this range was influenced by the need to prioritize the first pattern and trial and error. Nevertheless, no threshold was identified, and thus, a stable system could not be achieved.

The cross-talk can be categorized into four distinct levels, each with its distribution pattern, Fig. 3.

**Figure 3.**
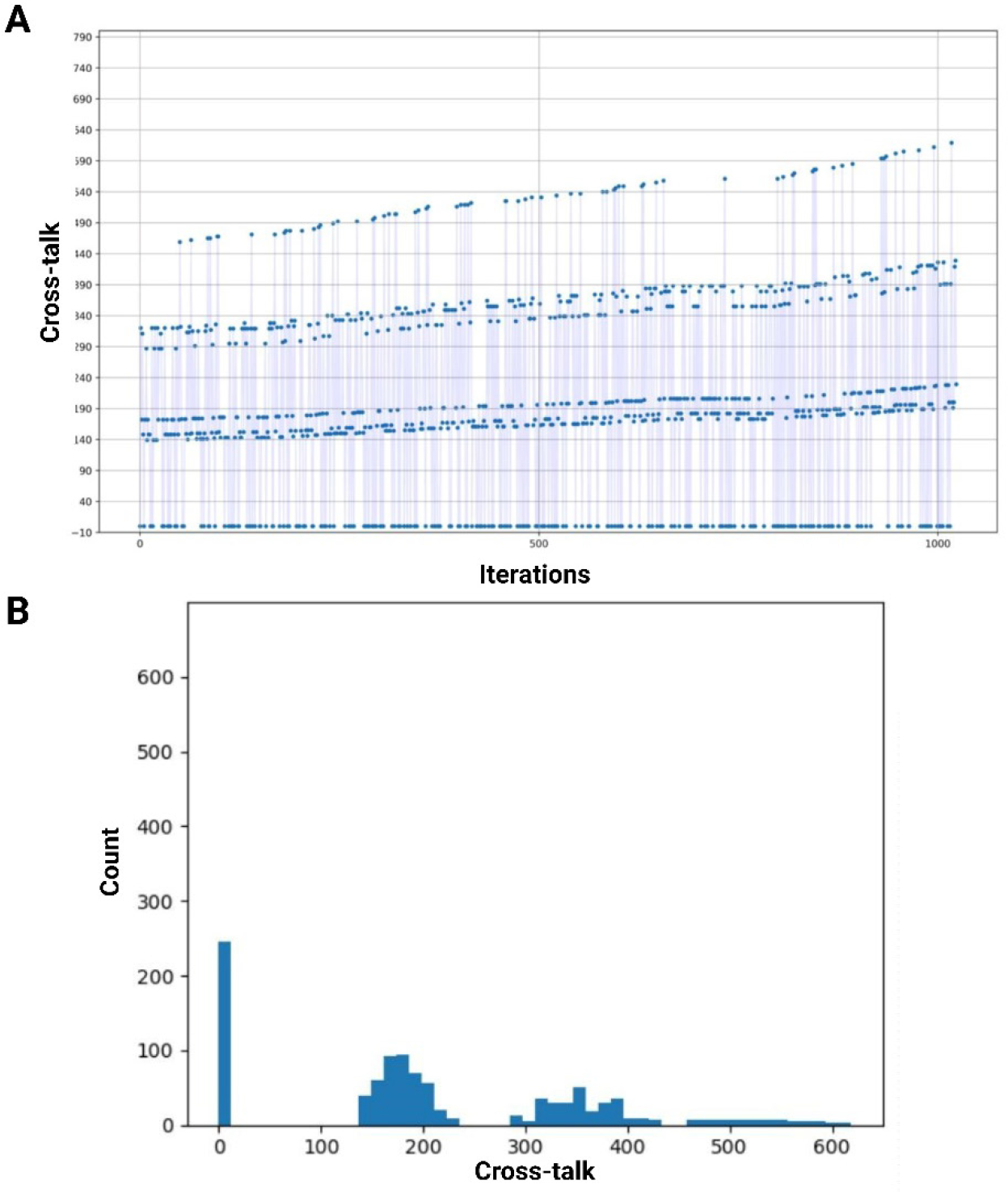
Illustrating the categorization of cross-talk into four distinct levels, each demonstrating its unique distribution pattern. Panel A depicts the iterations for the cross-talk, categorized into four distinct levels. Panel B shows a histogram displaying the frequency distribution of the cross-talk counts. The first level pertains to scenarios in which the j-th neuron does not encounter any form of cross-talk. The occurrence of the second level is characterized by the interference of only one pattern with the asynchronous update. The third level pertains to the interaction between two distinct patterns. The fourth level signifies the most unfavorable situation where all patterns intersect concurrently.

The first level relates to situations where the j-th neuron does not experience any cross-talk. The second level occurs when only one of the patterns interferes with the asynchronous update. The third level involves the interference of any two patterns. Finally, the fourth level represents the worst-case scenario where all patterns interfere simultaneously.

The changing network state is raising the average cross-talk for each scenario, which is another sign that the training pattern we fed into the system is unstable. A stable pattern would lead to a constant average cross-talk; however, some levels are more prevalent. More iterations had zero cross-talk than any other number, and all patterns contributed to cross-talk least often.

Our pattern distribution of ones and zeros interferes with asynchronous updates. Therefore, we reduced the p-value for our training patterns to *p* = *P*(σi = 1) = 0.2, resulting in half the average number of ones compared to the simulation before. The sparser distribution should reduce interference on average for all four levels except 0.

The simulation was thus performed for M = 4 patterns with the following square norm matrix:

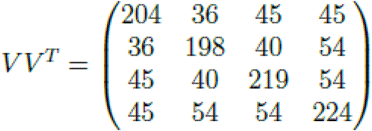

We biased for the first pattern and picked a threshold slightly less than the norm squared to detect the most robust pattern while muting all others. These patterns would be within the range of perfect thresholds if the training patterns were orthogonal or pseudo-orthogonal, Fig. 4.

**Figure 4.**
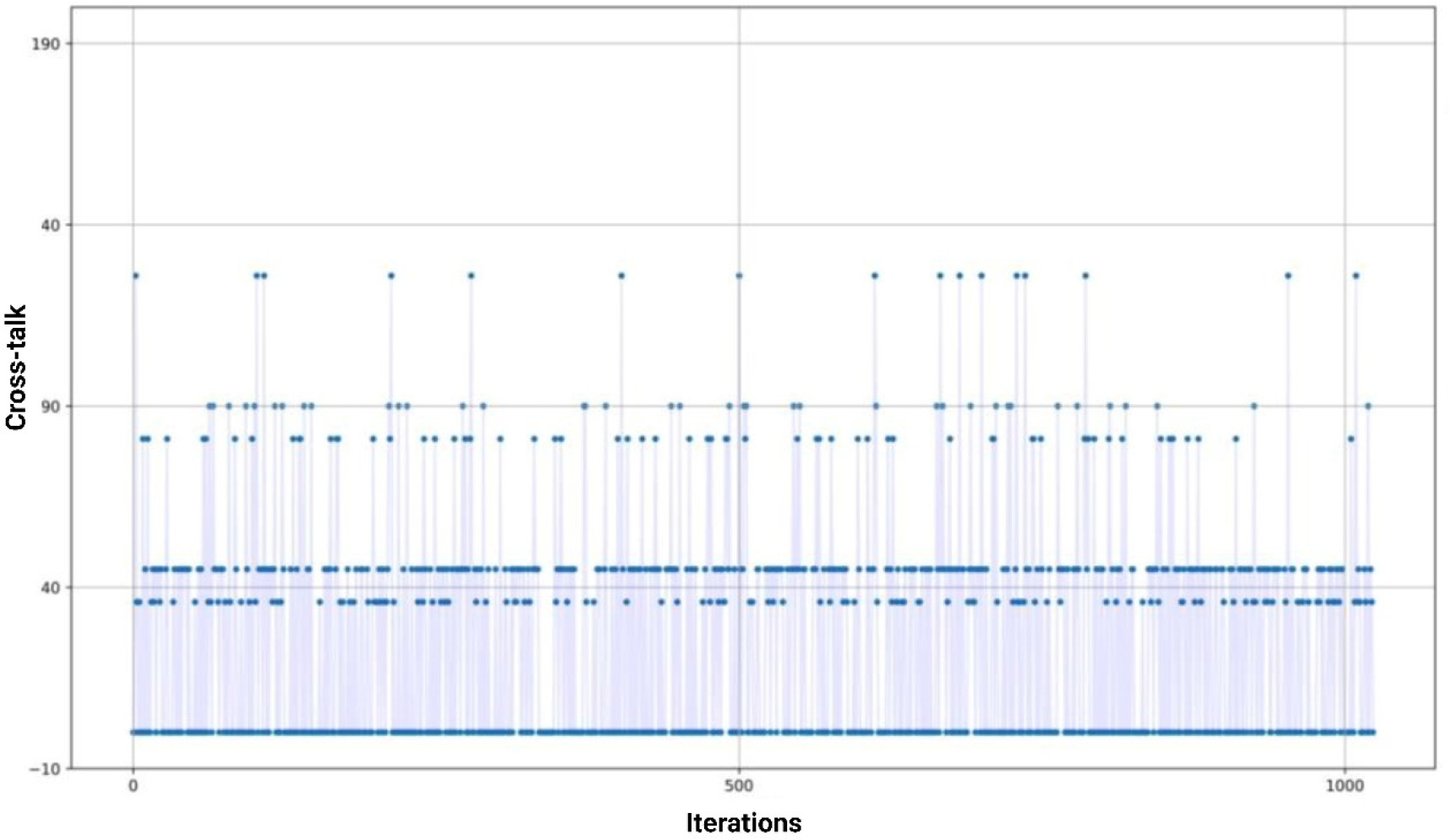
Depiction of the bias toward the first pattern and selecting a threshold slightly below the square of the average value to identify the most resilient pattern while suppressing all others. If the training patterns were orthogonal or pseudo-orthogonal, these patterns would fall within the range of optimal thresholds.

The stability range for this simulation was (126 <ϕ < 204), as the threshold mutes cross-talk and biases for the most robust pattern, the first pattern. We have improved stability by selecting training patterns with more sparsity in activation.

Simulations have been performed for M = 5, 6, 7, and 8 to determine how much a 20% distribution boosted memory capacity. We chose input patterns with the least squared norms from the training patterns for each simulation because stability for the entire network was ensured in our simulation when the least norm training pattern was stable.

Stability was always maintained for M = 5, even for small threshold ranges, between the squared norm of the input pattern (also the training pattern) and its dot product with the highest interfering pattern. However, stability was not ensured for M = 6 and above; as a result, we reduced the average distribution of ones per pattern to 10% of the total number of neurons.

A memory system with a limited capacity to store only five distinct patterns is not especially useful, particularly when considering emulating the mechanisms of actual cell assemblies. Consequently, further simulations were performed using a probability value of *p* = *P*(σi = 1) = 0.1 and starting from M = 5. This step was necessary to establish that HNN possesses adequate capacity for the subsequent phase of modeling creativity-based associations. The square norm matrix for the training patterns was determined as follows:

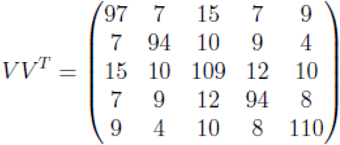

This matrix was stable, with a threshold in the range (92 < ϕ <60), which is still consistent with the threshold ranges that we previously indicated. Finally, simulations were performed for M = 6, 7, 8, 9, 10, 11, and 12. Stability was ensured up to M = 8 but not above; stability rarely occurred with M > 14.

### 2. Reconstructing Associative Chains

Gabora (2010) argued that content addressable memory requires appropriate sparsity in activating patterns across a cell assembly. A fully distributed memory would store all memories across all neurons, causing memory conflict and recall instability. In contrast, fully localized memory is only kept at designated locations; recall requires knowing the neuron (Gabora, 2010). Accordingly, Content-addressed memory operates effectively with sparse, distributed memory, and the brain’s density makes this possible. To gain meaningful capacity for our HNN model, we implemented certain modifications.

Returning to the updated rule description, we find approaches to improve the recollection of patterns. Equation 2.4 shows that the middle sum can be used as a weight for the kth pattern and its contribution to the network’s jth neuron update:

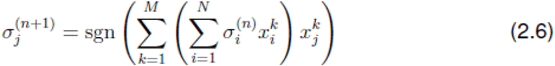

We rename this weight component as:

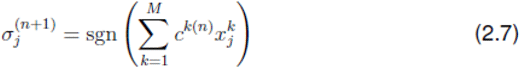

The weights term can be regarded as a parameter that varies in response to system state changes. We update as weight controls neuron cross-talk. One approach could involve adjusting our network architecture such that only the pattern with the highest weight is considered at each update. The procedure employed in this approach resembles our strategic approach in determining a threshold value that ensures the stability of all training patterns as system points. The weights matrix that was previously defined is not included in Equation 2.7. Although the current structure appears distinct from our original model, it is crucial to acknowledge that it represents an alternative perspective of HNN.

Instead of using a weights matrix, we use training patterns as weights in a two-layer network. The first layer represents the network state (input and output), while the second layer stores the 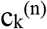 values, which are the inner product of the network state and each training pattern. The weights (training patterns) are the same from the state layer to the hidden layer and vice versa; thus, information is transferred up from the input to the hidden layer and down from the hidden to the output layer in each iteration. This view also shows how each pattern activates state layer neurons. We have reduced updating and converging the network to a competitive structure at the hidden layer whose winner should always be closest to the network’s present state.

After managing the “influence” values as a distinct layer, we employ an activation function on this hidden layer (we do not name them weights to avoid confusion with the recurrent network’s training patterns). Initially, the p-norm was used with a p-value of 1, which only normalizes the weights matrix within the range of 0 to 1. The largest influence would not be biased over the remainder of the noise. Normalization is formulated as the following:

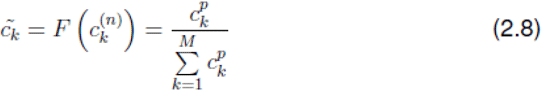

We observe significant storage memory capacity gains at p > 5. For p > 10, employing training patterns within a few striking distances of each other limits capacity since multiple patterns seem the same, making it impossible to recall the correct one. This is similar to how our brains consolidate all experiences of an idea rather than storing them separately. We can get nearly flawless recall by setting the sigmoid activation to 0.5 from a p-norm of ≥5. Our new update then has the following form:

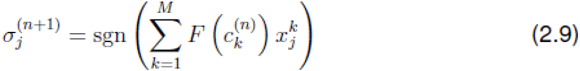

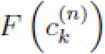 represents the activation of the influence/hidden layer, such as the p-norm or another suitable alternative.

This modified description of the HNN is similar to Krotov and Hopfield, which was considered a high-capacity discrete neural network derived from an update only minimizing an energy function for the whole system (Krotov & Hopfield, 2016).

They described an update with the form:

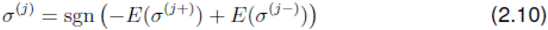

When the jth neuron of the present state is flipped, the energy of the system is E(σ^(j−)^), and when it is not flipped, it is E(σ^(j+)^). The activation pattern equation, σ = (σ_i_) ∈ {−1, +1}^N^, maintains that jth neuron activation only applies if it lowers system energy. The system’s energy was defined using an interaction function to ensure far-distant local minima and accelerate the convergence of the HNN model, which was subject to cross-talk and entrapment in spurious local minima. Similar to our modification, it biases for the nearest pattern and mutes cross-talk simultaneously.

This form of the energy function is defined as:

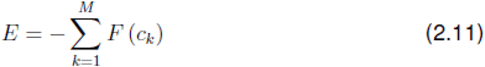

As previously described, ck indicates the effect of M training patterns on each j-th neuron update. Krotov and Hopfield defined an interaction function, F(z) = z^*p*^, which resembles the p-norm used to activate/normalize the hidden layer. Accordingly, the modifications provided resemble a feedforward structure, making them appropriate for deep neural networks. It is similar to the biological recurring structure to boost capacity by minimizing interference.

Ramsauer et al. (2021) introduced a continuous HNN model, which follows the Krotov and HNN modification and includes a new energy function for continuous-valued patterns. This modified HNN also utilizes an activation function and An energy function rule updated by the concave-convex approach, yielding a softmax function as the hidden layer activation function (Ramsauer et al., 2021). The energy function was based on Ramsauer et al. (2021) for patterns with continuous neuron activations; however, it is effective because we employed softmax activation in the third layer.

This activation is formulated as the following:

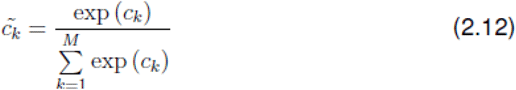

Our algorithm’s effect on the system’s state space navigation was demonstrated by mentoring the system’s energy at each iterative step, resulting in a figure that depicts the energy path taken over time (Fig. 5).

**Figure 5.**
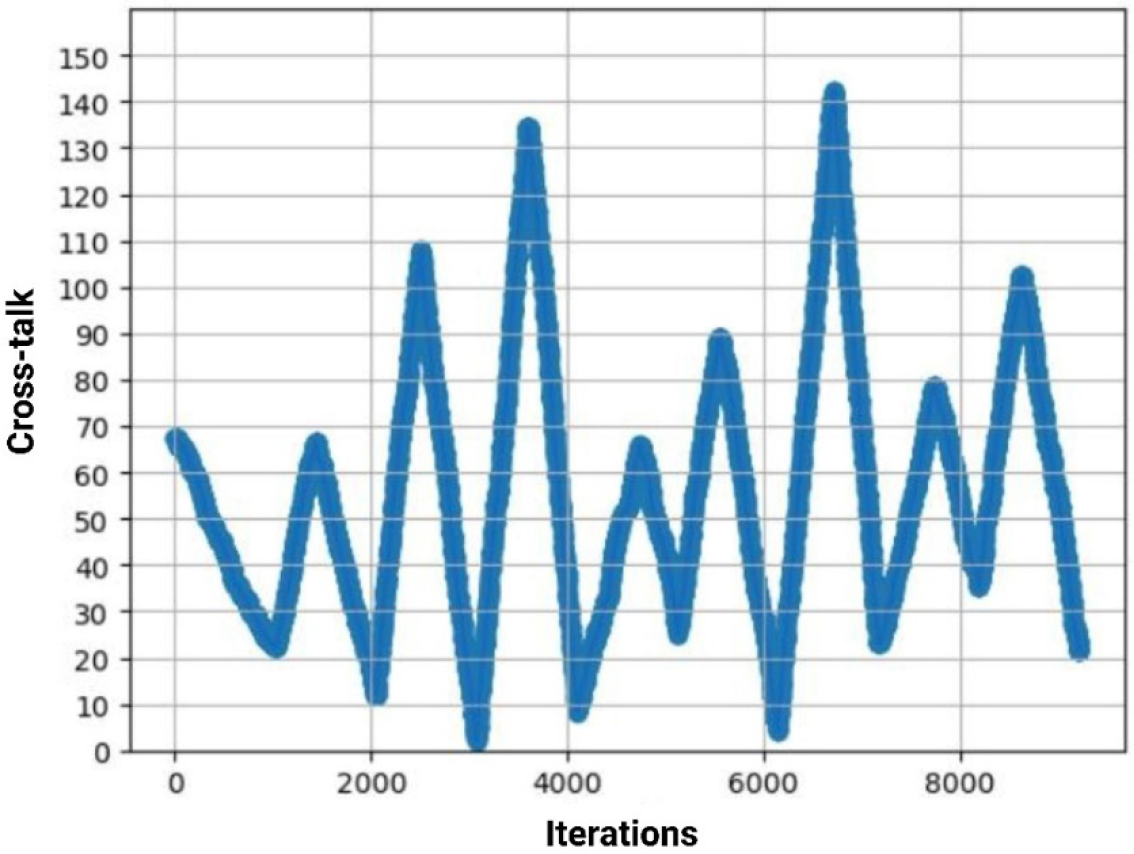
The energy path through time illustrates how the algorithm influences state space navigation and energy at each iteration.

Therefore, it transforms our hidden layer activations into a probability distribution, which is biased extensively for the node with the highest influence and exponentiates the capacity of the HNN model. Consequently, Semantic association-based creative thinking involves connecting previously unrelated ideas to develop a new path (Beaty et al., 2023; Kenett & Beaty, 2023; Li et al., 2021; Luchini et al., 2023) could be explained through this model.

## Results

Our main result shows how this new parameter affects convergence by acting as a weight parameter distribution of 1s and 0s over each learning pattern in the unipolar binary case layer, passing the value down to the hidden layer and completing one update loop (Fig. 6).

**Figure 6.**
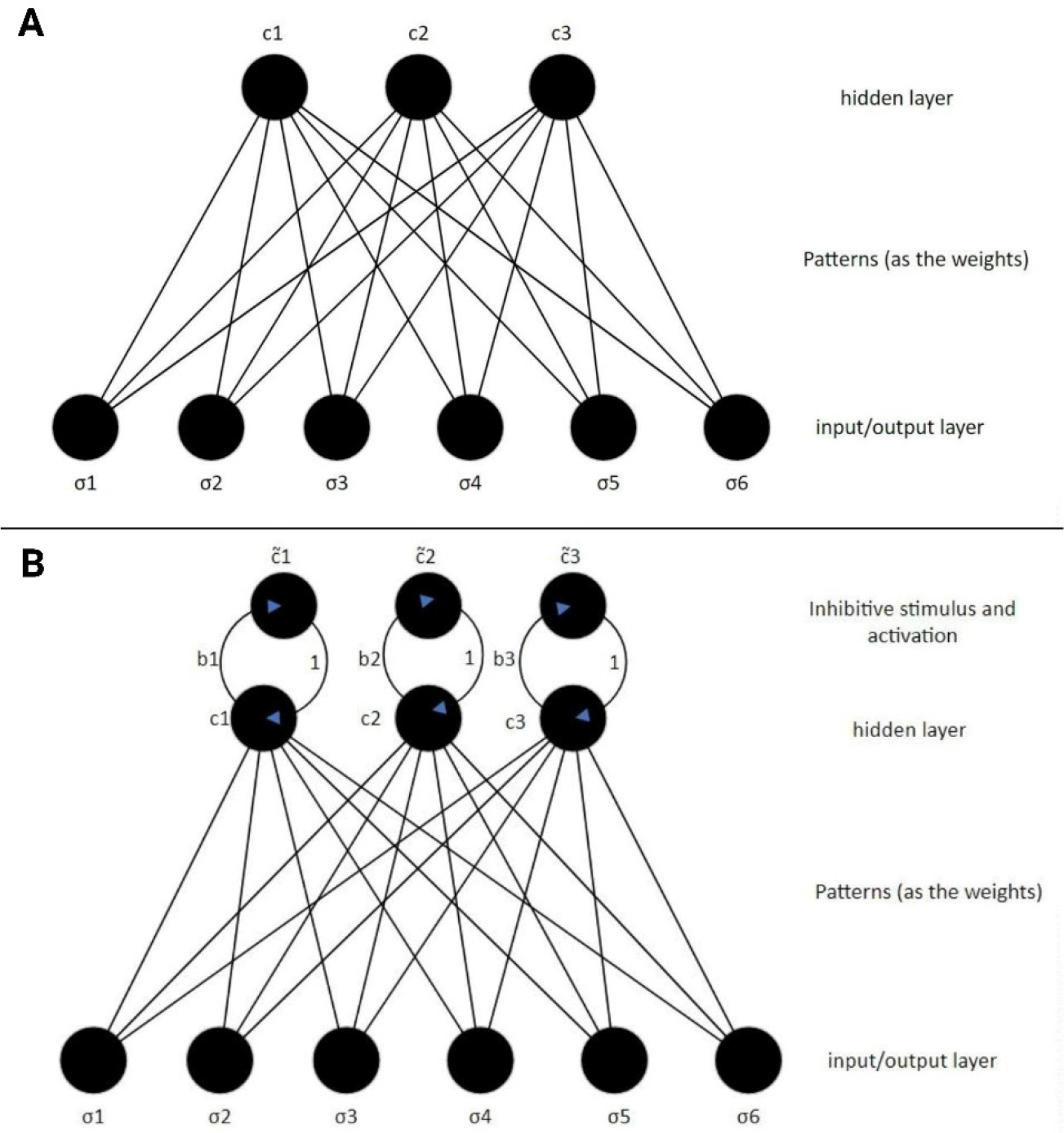
Demonstration of how adding a new parameter affects convergence by being a weight parameter distribution of 1s and 0s across each learning pattern in the unipolar binary layer, delivering the value to the hidden layer, and completing one update loop. Panels A and B show the two-layer HNN model with inhibitive stimulus and activation adjustments

Our algorithm treats inhibitor biases as a system hyperparameter. After adding the inhibiting stimulus, we have the modified equation:

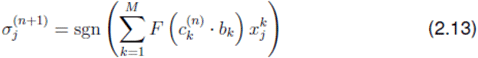

The hyperparameter can take values 0 ≤ b ≤ 1; we define the influence inhibition as a percentage value, where no inhibition corresponds to 100% of the influence being allowed, and complete inhibition corresponds to 0%.

The algorithm for iterating this hyperparameter defines that each whole network update corresponds to one set of inhibitors, which are set while updating the first neuron. The algorithm determines which hidden layer node has the most significant value, the concept closest to the input state, an unknown pattern stimulating the system. In the initial iteration, our system recalls the first link in our associative chain, the closest association. We initiate a new iteration loop to locate the second link, starting with the concept from the prior system update. The system retains the information on the nearest associated idea, the same concept recalled before when the largest hidden layer node is known. That hidden node is muted before the third layer is triggered. The concept’s effect has decreased; activation will choose the next closest associative choice, and neuron updates will converge until the following link is complete.

In the following major update, inhibitors reduced the previous two influences, allowing for the closest idea to the chain. This process is continued until a desired length of associative links is reached to diverge from prominent to hidden associations.

Once the network has an appropriate associative chain length, this process stops. Simulations showed this inhibitive connection process using softmax() as the third layer activation function.

To select stored concepts, M = K + nK, where K is the same random *p* = *P*(σi = 1) = 0.2, N = 1024 patterns were utilized, and nK is n randomly perturbed versions of K patterns with 100 hamming distances between them.

Therefore, this simulation used K = 3 patterns. We constructed n = 2 perturbed versions of each pattern, with the first using 100 random bits and the second using 200 bits from the prior perturbation. This procedure yielded M = 9 patterns with predetermined associations, enabling us to identify the final associative chain.

After introducing a random and unknown input into our network, the network was permitted to find the first associative link among the known M concepts that correlated best with the input. This determination was achieved after just one complete update, and the next link was formed by inhibiting this first result and the next one by inhibiting all previous recollections. The known patterns, ordered from most to least correlated with the initial unknown input, generated a unique M-length chain.

Our pattern selection reveals the following two links in the chain after the initial surprising one was formed. Even though the patterns’ correlation levels were semi-hard coded, the established chain was surprising given our lack of input state knowledge and the fact that these were randomly generated patterns. Thus, with a larger system with more neurons that know hundreds of thousands of concepts, many surprising associative chains are possible, giving the system surprising insight into the input stimulus based on what is already known. The more complex the chain, the greater the degree of unknown and surprising the results.

## Discussion and Concluding Remarks

We first attempted to establish a network with sufficient capacity to be scaled to large numbers of concepts, which is necessary to form meaningful associative links. After our first simulation experiments, we found the original HNN model unsuitable because it had almost no resilience to cross-talk from correlated concepts, another critical aspect of creative thinking-based semantic association. After examining the network topology, it turns out that it is a correlation factor, which was exploited to add a hidden layer and update based on correlations. Coupling the hidden layer with an activation function, such as a p-norm or softmax, was an effective biasing for the concept with the highest correlation with the input, exponentiating the network’s capacity. Our modification resembled the discrete and continuous current HMN (Hopfield, 1982; Krotov & Hopfield, 2016; Ramsauer et al., 2021). We created associative connections between previously unconnected concepts using HNN to establish a creativity-based semantic network conceptual framework (Fig. 7).

**Figure 7.**
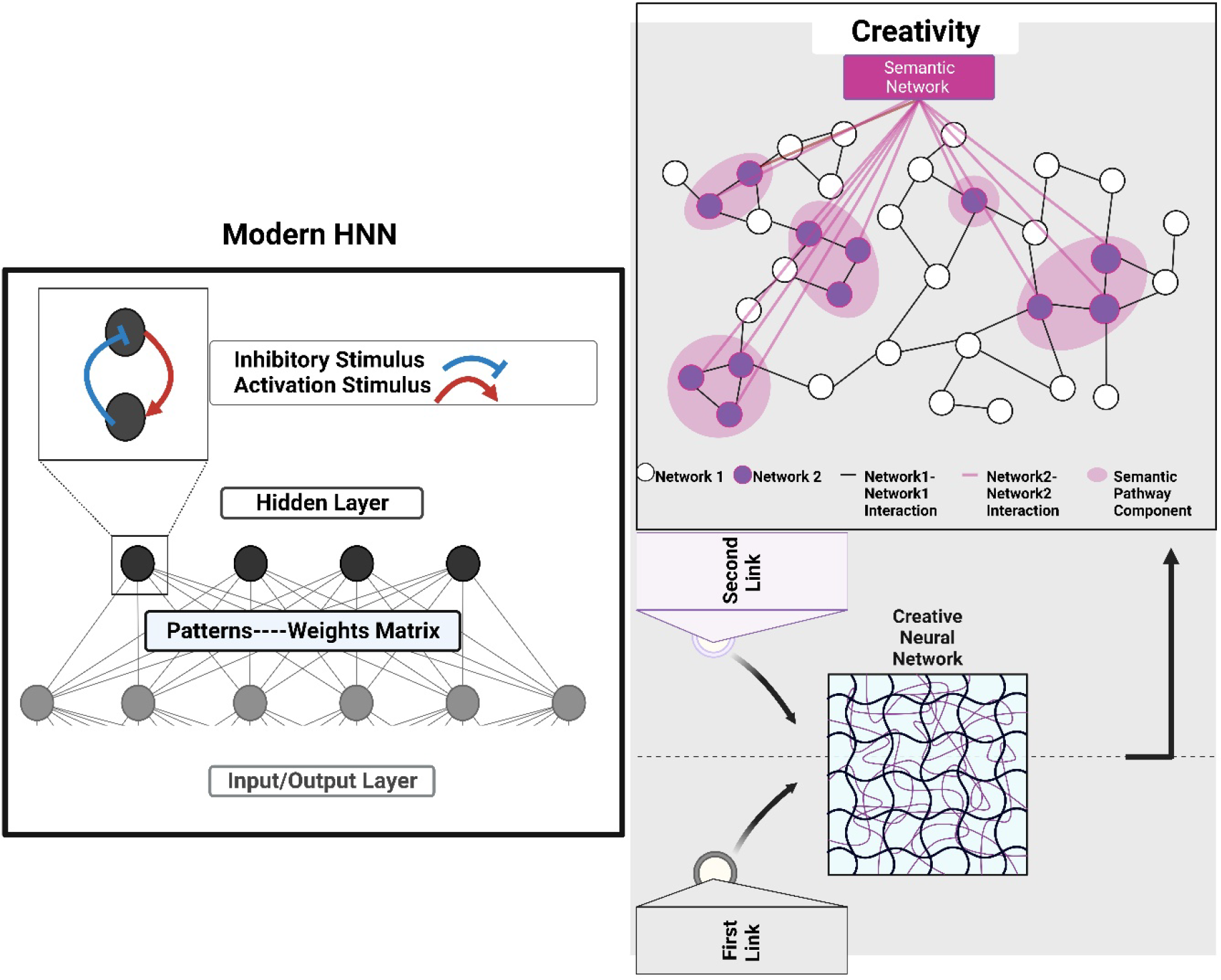
Summary of a hypothetical model of the simulation experiment outcome. Modern HNN is considered a suitable model to represent the associative chains of creativity rather than binary HNN. We created modern HNN as a two-layer model. The first layer comprises the neurons of the input, and the connections between the two layers are realized through the neurons of the patterns. The top layer contains the core of the updating process, which may be regarded as the “new weights.” The ‘new weights’ reflect how adding a new parameter affects convergence by being a weight parameter distribution of 1s and 0s across each learning pattern in the unipolar binary case layer, delivering the value to the hidden layer and completing one update loop. Accordingly, this resembles creativity’s first and second links, symbolized as a semantic association.

We developed a model that can store multiple patterns, showing concept associative linkages, which align with the definition of context focus and show how our brain relates concepts to link them together as they search for optimal solutions to creative problems (Gabora et al., 2010). To facilitate convergence, we introduced a new parameter in our network that inhibits the influence of previously learned concepts, “associative link”.

Using modern HNN, we identified two “mechanisms” controlling the context focus, shifting from analytical and cause-and-effect thinking to associative-based thinking. The first mechanism refers to the activation threshold of neurons, which acted as an on/off switch for our network. The second was the inhibition of stored concepts, which was similar to an on/off switch that guides the system when to search for associative links and when to stop.

Thus, inhibition could act as an additive inhibitory mechanism besides the activation function used in the state layer, whose threshold can be considered another on-off switch controlling whether the system can converge to an association. This finding agrees with empirical creativity studies that indicated the influence of inhibition in creative thinking (Benedek et al., 2012; Carson et al., 2003; Green & Williams, 1999; Khalil et al., 2020, 2023; Radel et al., 2015; Scibinetti et al., 2011). Our findings suggest that inhibition is essential for changing our focus from cause-and-effect stimulation (analog to analytical thinking) to correlations and associations (analog to creative thinking-based semantic associations).

The first and second links assisted us in creating associative links between previously unrelated concepts and further developing them computationally into associative chains connecting several concepts; for a better understanding of the concept, see Fig. 8.

**Figure 8.**
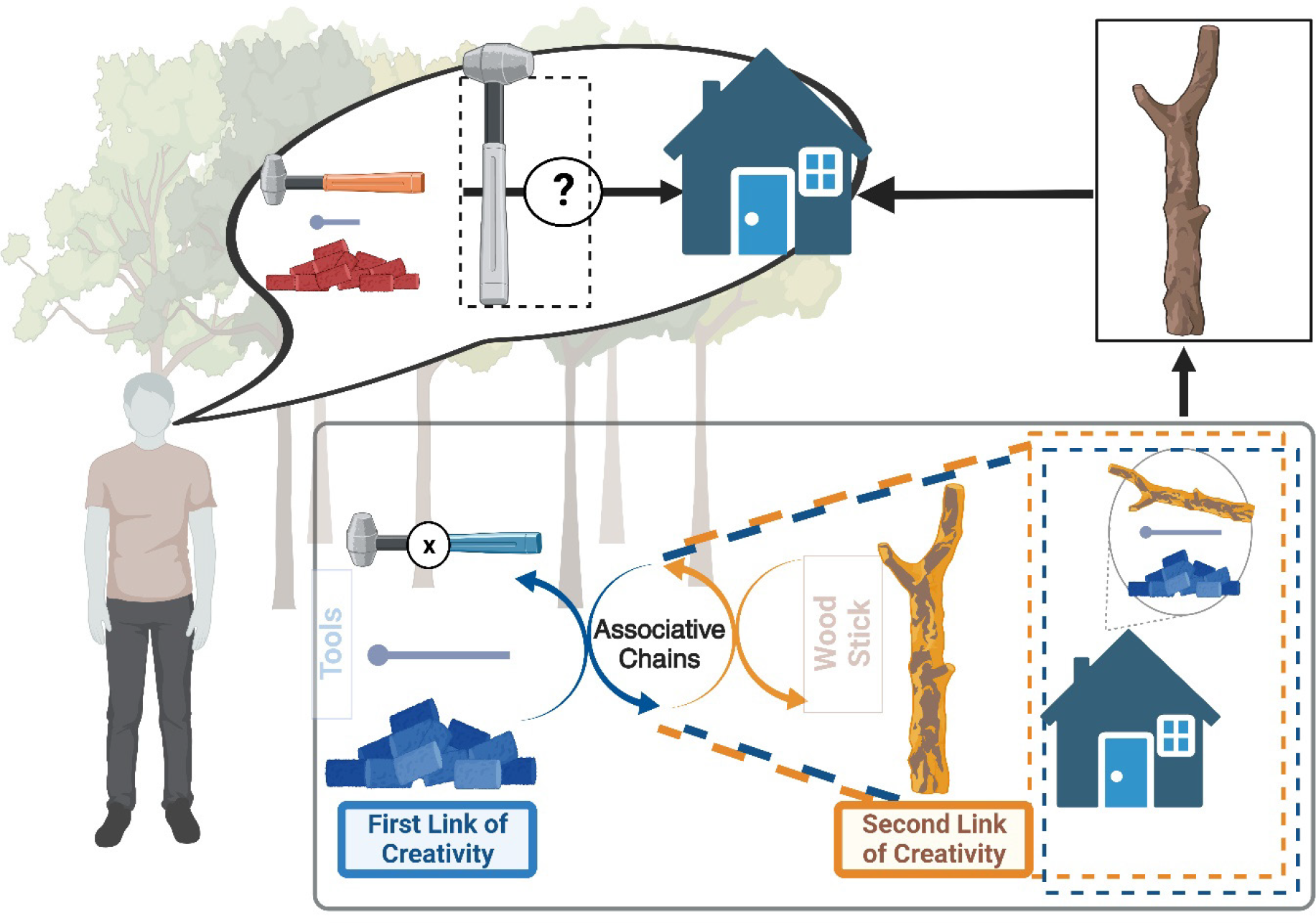
An example of how concepts are linked and how far our associative chain may extend without presumptions. A man in the forest builds a house, which is our example. This man has learned to build a house with hammers, nails, walls, and rocks. If the man is shown a nail and a wall indicating driving a nail into the wall, he may think of the hammer first. However, if he is in the forest and has left his hammer at home, he may think of something else that is more directly associated with the hammer, such as a rock. Because he could deviate from the hammer, he allowed himself to form an association that led to a solution. The inhibition can be considered a mechanics switch that directs the system to engage in associative thinking, which is the divergence from evident to obscure relations; this allowed him to solve his problem creatively.

In conclusion, we successfully demonstrated the first neurocomputational framework model for creativity-based semantic associations. This model was built using both a simple, binary HNN and a modern, two-layer HNN (Fig. 7). Our model suggests that we do not have to search the whole network and compare our input with all the patterns to find a solution; it helps us retrieve the memorized items seamlessly using inhibition as an analog to the hyperparameter. Several creativity studies have highlighted the role of inhibitory control on creative thinking (Benedek et al., 2012; Carson et al., 2003; Green & Williams, 1999; Khalil et al., 2020, 2023; Radel et al., 2015; Scibinetti et al., 2011).

Therefore, an associative link is created when a group of neurons (i.e., neural cliques; Wang et al., 2003) fire up together while being part of both concepts we are connecting; this allows us to find a solution that would not be an oblivious choice, but that solves our problem. These connections are still considered obvious and are activated by slightly different groups of neurons, known as neural cliques (Wang et al., 2003). This association could be analogous to creative thinking based on semantic associations, allowing us to search on the lower levels of memories to find which concepts relate even on a micro-level to our situation (Beaty et al., 2023; Kenett & Beaty, 2023; Li et al., 2021; Luchini et al., 2023). The patterns and how we store and reconstruct them align with the biological processes, as human memory is also distributed, coarse-coded, and content-addressable (Beaty et al., 2023; Benedek et al., 2023; Gerver et al., 2023; McEliece et al., 1987). Another potential explanation of our findings is that our neurons take a step back from the contextual focus and find alternatives associated with seemingly unrelated concepts when analytical thinking is insufficient, using inhibition as an analog to the hyperparameter.

This preliminary computational neural network model has numerous possibilities for investigation, as we can find creative ways to connect concepts and see how far our chain can reach without imposing a predetermined limit on the extent of these connections. Throughout this paper, the story of the man building a house in the woods is used as a visual aid to explain the dynamic application of our HNN (Fig. 8). We demonstrated how this neural model can effectively store associative and content addressable memories and access them in an associative manner, given an input excitation, forming what we call associative links that are essential in the creative problem-solving process.

### Future Direction and Limitations

Our findings are encouraging for the subsequent progress we can implement for model development, especially regarding the shift in the neurons from discrete to continuous activation, allowing us to have a greater variety of concepts based on the shades in the activations. It would also allow us to apply inhibition with more dexterity on the concept’s features, potentially making even more surprising and context-specific associative links. Krotov and Hopfield (2016) highlighted the duality of their discrete modern HNN and the feedforward structures of deep neural networks. Therefore, one can also imagine how our associative links algorithm may be integrated into a larger neural network and provide a mechanism for problem-solving through association. The potential for a more comprehensive creative framework is fascinating, but several remaining questions remain open.

Several fascinating unresolved explorations necessitate attention in the coming study endeavor; presented below is a representative example. Memory representations of visual, tactual, and auditory perceptions constitute the word “bell”. These memory images show the bell’s essential characteristic features. Therefore, it would be ideal for testing the framework on patterns representative of real-life concepts (i.e., create a dataset for them and define a set of multiple features that should be defined for each of them (e.g., human/animal/object, weight, height, color, etc.) as a next step, daily creativity (Richards, 1993; Runco & Bahleda, 1986). While it can present empirical challenges, developing an algorithm to process a given idea as input is feasible. Data can be collected and subsequently transformed into an identifiable pattern by providing questions to the individual regarding the concept’s features. The concept of a feature can be comprehended as a cluster of neurons, such as the initial 10 neurons within each pattern. Its definition depends on the activated neurons’ arrangement, where a neuron’s firing in the second position within a weight feature signifies a light object. At the same time, a value of 0 indicates the absence of weight relevance.

Expanding this network in terms of large-scale neural networks by testing this theoretical framework on concepts modeled by real-life concepts accordingly, the network’s creativity-based semantic associations will be more straightforward to observe effectively. Taking it a step further, we could redefine how associative links get created by iterating solely on the defined features and suppressing certain ones.

## Notes

### Competing Interest Statement

The authors have declared no competing interest.

